# Ecological opportunity and the onset of polyploid niche expansion waves

**DOI:** 10.1101/2025.07.11.664136

**Authors:** Felipe Kauai, Yves Van de Peer, Dries Bonte

## Abstract

Polyploidy, the presence of more than two sets of chromosomes, has evolved many times across the tree of life, yet we still do not know why some polyploid lineages persist while most go extinct. The establishment of polyploid populations is often reported to be associated with harsh environmental conditions, and stress tolerance in particular, which have led to the widespread view that polyploidy-specific niche requirements are key to their persistence at ecological and evolutionary timescales. Here, we reevaluate this perspective through a classical mathematical model of polyploid establishment, for which we provide new analytical and numerical results, along with an empirical case study. We show that simple eco-evolutionary processes at the margins of diploid range-expansion waves, more specifically ecological drift and dispersal limitation, can be sufficient to allow polyploid populations to carve out their own space, without any *a priori* adaptive advantage over their diploid ancestors. Our modelling effort reveals three key insights. First, polyploids most readily gain a foothold at the low-density front of a diploid range expansion wave, where ecological drift is strongest. Second, limited dispersal accelerates spatial clustering of polyploid organisms through assortative mating. Third, once spatially segregated, diploid and polyploid populations experience different environments, so natural selection can drive niche divergence. We illustrate these interconnected principles by simulating the phylogeographic history of a well-documented autopolyploid complex of *Neobatrachus* Australian burrowing frogs. Altogether, our results provide a neutral baseline against which the ecological consequences of polyploidization can be readily detected and inferred within natural populations.

**Significance statement:** Polyploidy, having multiple copies of the entire set of chromosomes, underpins major innovations in both animals and plants and fuels biodiversity. Yet, why some polyploid lineages are successful while others fail remains unclear. Our study shows that simple eco-evolutionary forces, specifically ecological drift at the edge of an expanding range and limited dispersal, can enable polyploids to establish and become geographically separated from their diploid relatives, even without any inherent adaptive benefits. This spatial segregation then exposes each cytotype to distinct environments, setting the stage for natural selection to operate. Our neutral theoretical framework helps biologists disentangle when polyploidy itself drives adaptation and when success is just a matter of being in the right place at the right time.

## Introduction

Polyploidization, the addition of one or more full sets of chromosomes to a diploid genome, may follow hybridization between species (allopolyploidy), or be the direct consequence of genome duplication within a single species (autopolyploidy). Polyploidy, and especially recent polyploidy, is prominent in flowering plants (1, 2) but has also been observed across many vertebrate groups (3) such as amphibia (4) and ray-finned fishes (5, 6). It has the potential to unlock novel genetic configurations that fuel evolution and promote the spatial and temporal diversification of life (1, 2, 7, 8). However, polyploidy is (generally) considered an unstable state. Genomic instability, genome loss, and overall fitness costs often lead to the systematic removal of such organisms from ecosystems over either ecological or geological timescales (9–12). Nonetheless, at any instance of time, cryptic polyploids or even entire populations thereof can be found across a wide range of taxa. How polyploids invade diploid populations and establish self-sustaining populations is an outstanding unresolved issue that constrains a more complete understanding of mixed-ploidy ecosystems.

In the presence of fitness costs, coexistence of related species (or cytotypes) is known to be facilitated by stabilizing ecological processes that lead to reduced competition, such as niche divergence (13). The assertion that polyploids reach successful establishment by the evolution of niche shifts away from the ecological niche of their diploid ancestors is a direct corollary of this observation (14–17). Early attempts to model the dynamics of polyploid populations had already suggested that disparate ecological requirements are necessary for polyploid establishment (9), but the mechanisms underlying niche differentiation between cytotypes have remained controversial. More recently, the emergence of niche divergence among cytotypes has been linked to a potential correlation between polyploidy and stress (2, 8), which may be the consequence of an expanded genetic and phenotypic diversity seemingly associated with genome duplications (18, 19). The success of polyploids has thus been ascribed to their ability to occupy and spread over environmental conditions that are stressful relative to diploid ancestors. Empirical data to support this hypothesis, however, are often inconclusive. Instances of niche contraction, expansion and differentiation have all been documented across many study systems (4, 15, 20–23). Moreover, current evidence suggests that niche differentiation is more consistent when analyzed in allopolyploids (15), whose ecological and evolutionary dynamics may be the consequence of the joint action of genome duplications with the merging of disparate genetic backgrounds. A focus on autopolyploidy is therefore needed to unravel the strict relationship between the whole genome doubling process and the eco-evolutionary dynamics of polyploid establishment.

Most models of polyploid establishment assume an autopolyploid origin, where syngamy of unreduced (diploid) gametes produced by diploid organisms leads to the emergence of a tetraploid individual (9, 24–28). In natural populations, average rates of unreduced gametes are very low, on the order of 10^−3^, with occasional pronounced variations (24, 29, 30). As a result, the syngamy of unreduced gametes is conditioned on the joint probability of independent and rare events. Given the strongly reduced viability of any haploid-diploid gamete combination, any establishment of a polyploid population must overcome strong positive density-dependent selection. In population genetics, the probability of fixation of a rare neutral allele can, however, be enhanced in small populations due to high genetic drift. For instance, low-frequency alleles may spread over large areas by surfing on the edge of range expansion waves, where population densities are small, competition low and drift large (31). Additionally, large intraspecific relative to interspecific competition resulting from, e.g., spatial sorting, may equalize fitness differences among species, and therefore promote coexistence of different species (32, 33). These observations suggest that a (pre)adaptationist perspective may not be required to understand the emergence and stability of mixed-ploidy populations.

Thus far, most theoretical work in the field of polyploidy has put focus on the rate of unreduced gametes produced by diploid cytotypes that is required for polyploid invasion. More recently, simulation-based studies have also been developed to study the effect of more complex aspects on the dynamics of populations with different ploidy levels, such as life history traits, demography and spatial structure (11, 26, 28, 34–37). However, these simulations display varying levels of complexity and substantially differ in their most elementary assumptions, which eventually blur their mutual connections and makes it hard to understand how they inform the theoretical understanding obtained from general solutions of analytical models. In fact, while computer simulations can serve as *in-silico* experimental systems, their role in extending basic theory is limited (38). Currently, complex assumptions used in simulations for the correlation between genotype and phenotype in polyploids lack a robust experimental basis and, as a result, claims about polyploid niche evolution with an underlying adaptive narrative remain controversial (19, 35, 39, 40).

To overcome bias from specific phenotype-genotype mapping and differential fitnessassumptions, we advocate that a thorough understanding of the establishment of polyploid populations necessitates a neutral ecological basis. A neutral, unbiased, model of polyploid establishment will then provide the ground state against which fitness differences that are ploidy-specific can be detected as potentially relevant ecological components. To this end, here we investigate the simplest model for the establishment of autotetraploids in a diploid population, as first considered by Felber (24), whose analytical results have only recently been obtained (27). Instead of considering only diploids and autotetraploids, we now also include triploid organisms of reduced fitness with variable production of gamete ploidy levels, more closely approximating the available empirical observations. We provide an analytical solution for a special case of the model and use heuristics and computer simulations for intractable results. Individual-based computer simulations are built to study the dynamical properties of the mathematical model in spatially concentrated populations *in-silico*. To illustrate our results, we finally use our model to simulate the phylogeographic history of *Neobatrachus* burrowing frogs, a well-described polyploid complex in Australia, by including a fitness map and mutation rates. Altogether, our results show that niche differentiation might evolve as a consequence of geographical differentiation and gradients in population densities, and hence, that differences in adaptive potential between cytotypes are not strictly necessary to explain cytotype dynamics. This stochastic and spatial perspective is put forward as a driver of polyploid niche evolution, independent of, or as a complement to, intrinsic adaptive processes that might originate from genotype-phenotype-fitness associations in polyploids.

## Results

### Cytotype dynamics in panmictic populations

We start by analyzing a simple, yet foundational, model of cytotype dynamics in a panmictic population. Instead of considering directly the time-evolution of cytotype frequencies, we study how proportions of gametes with different ploidy levels change over discrete time units within an initial diploid population with non-overlapping generations. Let *x*_*t*_, *y*_*t*_ and *z*_*t*_ denote the frequencies of haploid, diploid and triploid gametes in a large population of sexually reproducing individuals at time *t*. Then, random syngamy of gametes produces cytotypes whose frequencies are obtained by the expansion of the binomial (*x*_*t*_ + *y*_*t*_ + *z*_*t*_)^2^. Assuming all ploidies above 4 in offspring are unviable, the frequencies of gamete types at time *t* + 1 are given by the following coupled non-linear iterated maps:

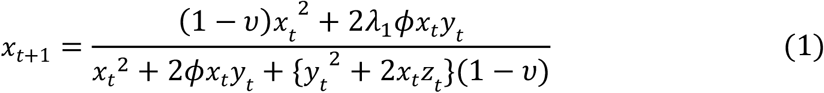

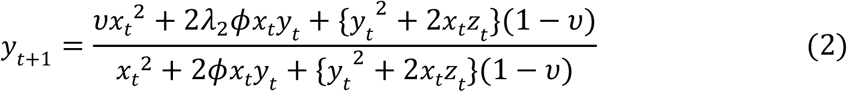

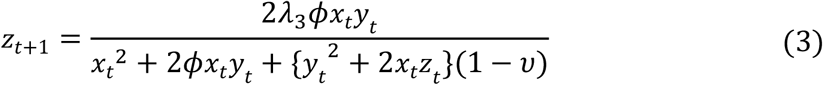

where *υ* denotes the rate of unreduced (diploid) gametes produced by diploid cytotypes (*x*_*t*_^2^), and *ϕ* the relative contribution of triploids (2*x*_*t*_*y*_*t*_) to the gamete pool, i.e., triploid fitness. The parameters *λ*_1_, *λ*_2_ and *λ*_3_ control the rate of haploid, diploid and triploid gametes, respectively, produced by triploid cytotypes. As with diploids, we assume tetraploids (*y*_*t*_^2^ + *x*_*t*_*z*_*t*_) produce unreduced (tetraploid) gametes with rate *υ*, which are neglected since syngamy events involving tetraploid gametes invariably result in offspring with ploidy above 4. Finally, the denominators of Equations (1 − 3) guarantee that *x*_*t*_, *y*_*t*_ and *z*_*t*_ are proper frequencies, i.e., contained between 0 and 1.

To approximate empirical observations of gamete types produced by triploid cytotypes (29), we first assumed *λ*_1_ = *λ*_2_ = 0.25 and *λ*_3_ = 0.50. That is, 50% of successful meiotic events in triploids produce unreduced (triploid) gametes, whereas the remaining 50% is equally distributed between haploid and diploid gametes. In this case, the system (1 − 3) cannot be solved analytically for the equilibrium points. To find the steady state of gamete frequencies with initial conditions *x*_0_ = 1, *y*_0_ = *z*_0_ = 0, we used a simple heuristic which is described in *SI Appendix Algorithm S1*. In figure 1A, we show the resulting cytotype equilibrium frequencies as a function of *υ* parameterized by *ϕ*. Notably, when triploids are unviable (*ϕ* = 0), the magnitude of *υ* at which tetraploids take over the system, referred to as *υ*_*max*_, was numerically found to be 0.20. We confirmed this result by considering a special case of the system (1 − 3) when *ϕ* = 0. Here, *z*_*t*_ = 0 ∀ *t* and *y*_*t*_ = 1 − *x*_*t*_. At equilibrium *x*_*t*+1_ = *x*_*t*_ = *x* and Equation 1 must thus satisfy:

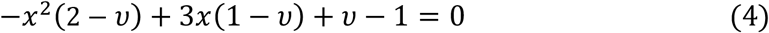

**Figure 1.**
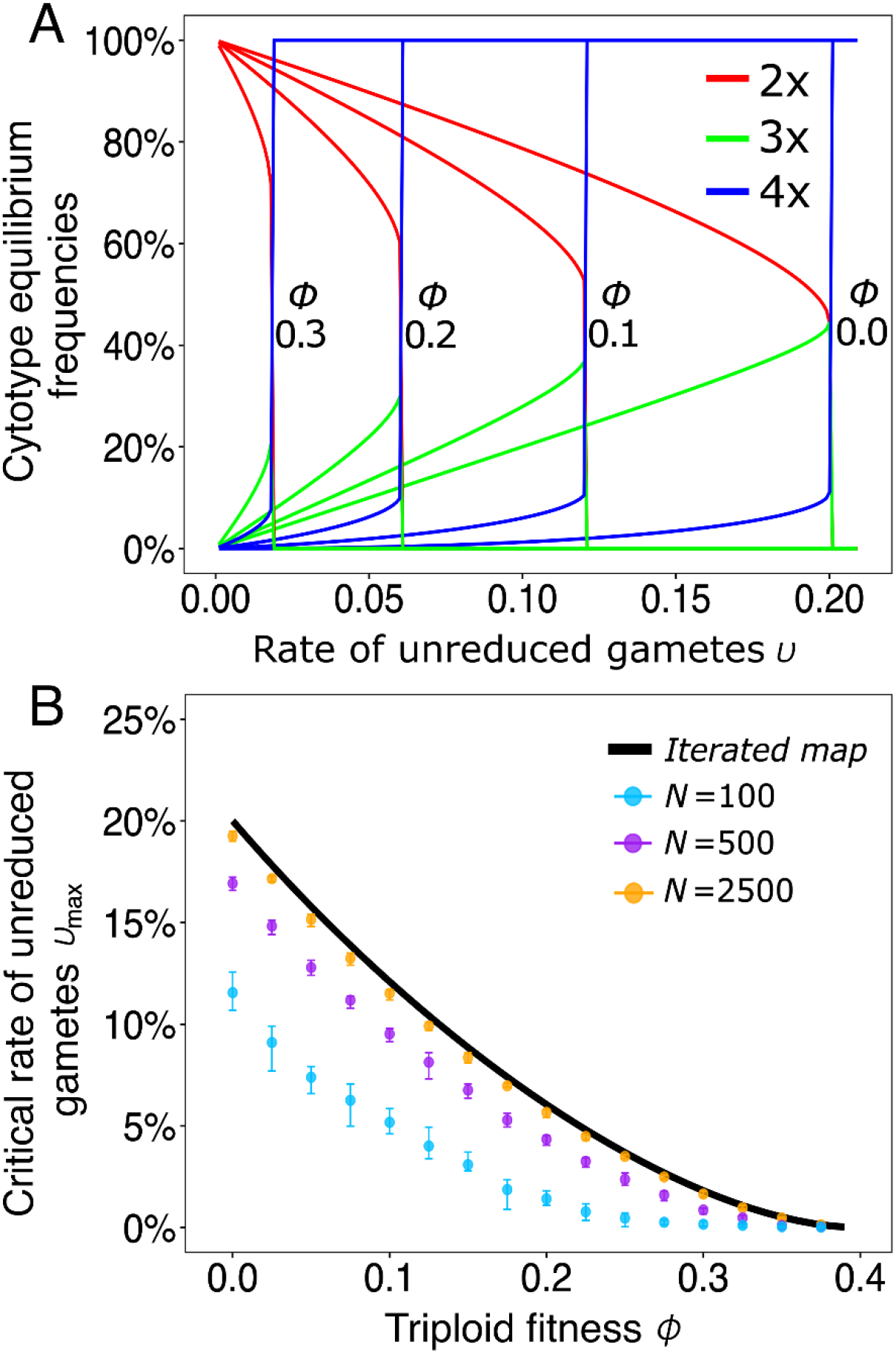
Equilibrium frequencies and conditions for polyploid invasion in a panmictic population. In Panel A) the cytotype equilibrium frequencies, obtained from the expansion of (*x*_*t*_ + *y*_*t*_ + *z*_*t*_)^2^, are plotted as a function of the rate of unreduced (diploid) gametes by diploid cytotypes *υ*. We factored them by triploid fitness *ϕ*. Thus, for each *ϕ* we show the converging equilibrium frequencies in the iterated map (1 − 3). Red represents diploids, green triploids and blue tetraploids. In B) the critical rate of unreduced (diploid) gametes, *υ*_*max*_, is plotted as a function of *ϕ*. Here, *υ*_*max*_ was obtained heuristically (see main text) for each *ϕ* in incremental steps of 0.001. The results from the iterated map are plotted in a solid black line. Average values from individual-based simulations are shown as closed circles for different population sizes (N). Whiskers represent ± 1 standard deviation from the mean.

Equation 4 describes a downward parabola with two fixed points (one unstable and another one stable) that depend explicitly on *υ* (see *SI Appendix Figure S1*). The magnitude of unreduced gametes *υ*_*max*_ represents the bifurcation point where both fixed points coalesce and are annihilated. Then, for every initial condition *x*_0_ the system always converges to 0, i.e., diploid gametes are the only gamete types in the system and therefore only tetraploid cytotypes exist. To find *υ*_*max*_ we first find the intersection of (4) with its first derivative:

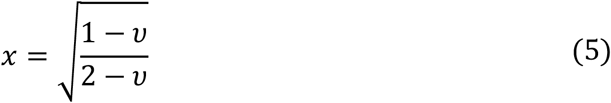

The bifurcation occurs when Equation (4) is satisfied given the constraint (5). Thus, substituting (5) into (4) and solving for *υ* we find that *υ*_*max*_ = 1/5, which confirms the result obtained heuristically. For the empirically motivated case (*λ*_1_ = *λ*_2_ = 0.25 and *λ*_3_ = 0.50) we computed *υ*_*max*_ heuristically across all *ϕ*. Remarkably, the rate of unreduced gametes produced by diploid cytotypes that is necessary for tetraploids to completely invade the system decreases approximately exponentially as a function of triploid fitness. In other words, for each unit increase in triploid fitness the frequency of meiotic errors during diploid gametogenesis necessary for tetraploids to completely invade the system reduces approximately exponentially. Moreover, individual-based simulations show that smaller population sizes, and thus, larger magnitudes of drift, significantly lower the invasion threshold *υ*_*max*_ (figure 1B) – a result similar to the increased fixation probability of an allele in small populations. Finally, to account for the variability often found in the gamete types produced by triploid cytotypes (26, 29, 41, 42), in *SI Appendix Figure S2* we highlight the effects of variability of *λ*_1_, *λ*_2_ and *λ*_3_ on *υ*_*max*_.

### Polyploid establishment in spatially structured populations

Now, we consider the same model as in the system (1 − 3), but assign two-dimensional spatial coordinates to every individual of a finite population of size *N* which inhabits a square lattice of size *L* = 100. We introduce an *extended* Moore’s neighborhood, R, which constrains mating to individuals that are within at most R grid cells from one another. For every generation *T* the system randomly selects a *seeker* and a corresponding partner among all of its neighbors within distance R. The neighborhood size, i.e., the number of grids cells around the seeker, is simply (2R − 1)^2^. Both individuals undergo meiosis according to the same rules of Equations (1 − 3), gametes fuse and the resulting offspring is randomly dispersed within the neighborhood of the seeker. The process is repeated *N* times for every generation, i.e., every individual has the same probability of successfully reproducing at every generation. An example of the resulting dynamics is given in figure 2A.

**Figure 2.**
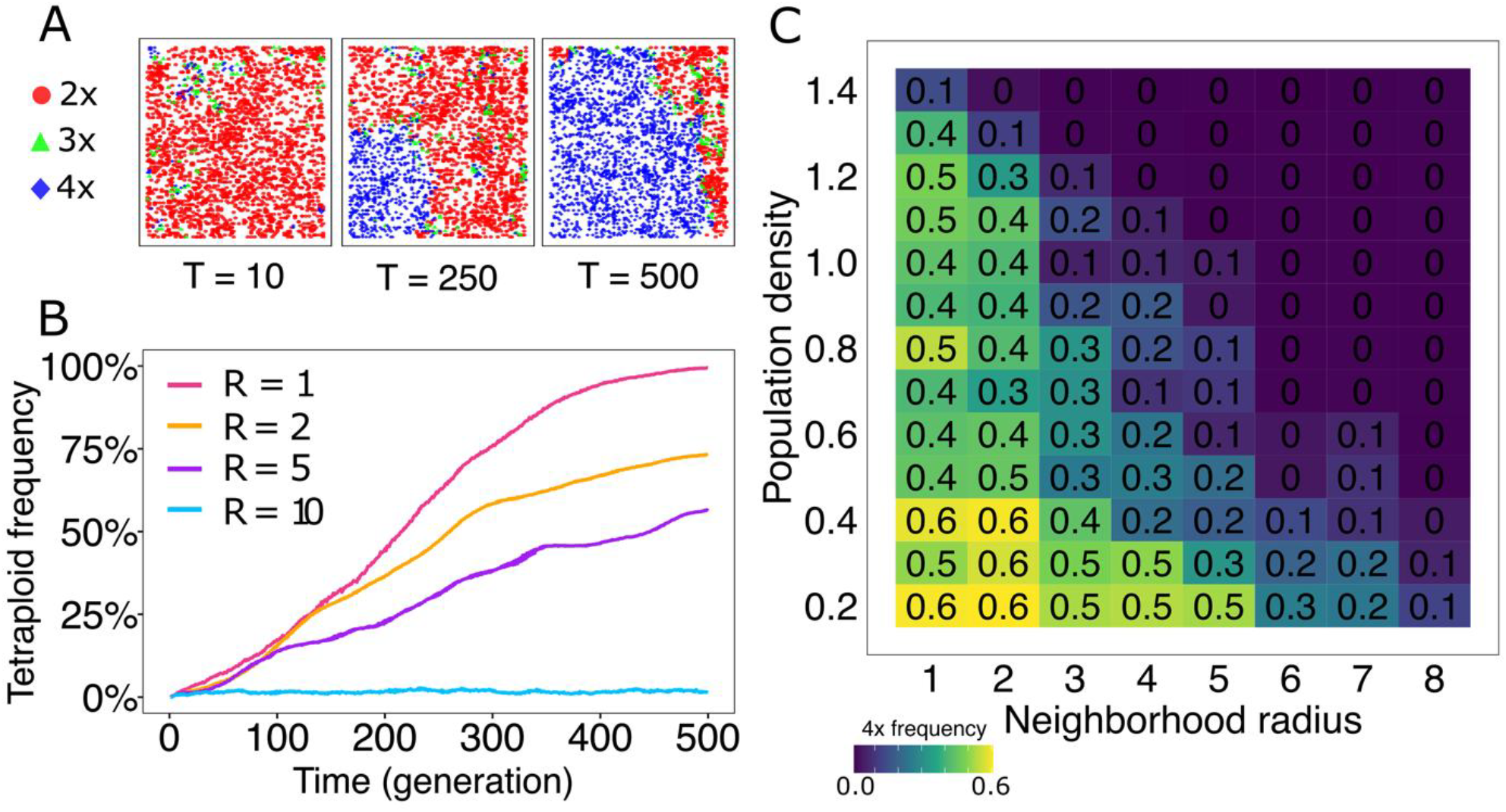
The effects of mating ranges and population densities on the spatial dynamics of mixed-ploidy systems. As an example, in A) we show three snapshots at generations T = 0, T = 250 and T = 500 of a single instance of the spatially explicit individual-based simulation of the iterated map defined by equations (1 − 3). In B) the frequency of tetraploids is plotted as a function of time factored by neighborhood radii *R*. In C) we plot the frequency of tetraploids (color gradient) at generation T = 250 as a function of population density (homogeneously distributed over the lattice) and neighborhood radii. Clearly, the frequency of tetraploids increases as both population density and neighborhood radius decrease. Essentially, this tile plot measures the speed with which tetraploids invade the system. Parameters used are *υ* = 10^−3^ and *ϕ* = 0.3.

As expected from previous theoretical and empirical work, a sexually reproducing polyploid population will cluster in space during establishment due to dispersal constraints (11, 34, 43). That is, the accumulation of polyploids in a certain spatial region will facilitate their reproductive success locally and thus initiate a wave of spatial spread. These spatial points of polyploidy accumulation are expected to occur primarily at the corners and edges of a two-dimensional landscape. Indeed, consider a homogeneously distributed population on a two-dimensional square continuous landscape of size *L*^2^. If the mating range of an individual is *R* ≪ *L*, then a seeker at the center of the landscape will have 2*ρπR*^2^ mating partners, where *ρ* is the population density. In contrast, there are only 0.5*ρπR*^2^ mating partners available for a seeker that is located right at the corner of the landscape, since the disc sector is exactly 1/4 of the disc in the center of the landscape. We have shown earlier that reduced population sizes facilitate polyploid takeover by reducing *υ*_*max*_. It is thus natural to consistently observe the focus of polyploid spatial spread at the corners and edges of a landscape. By the same token, smaller mating ranges will facilitate polyploid establishment by constraining the number of available mates to a given seeker (figure 2B), whose mating opportunities will decrease with the square of its mating range. Naturally, when *R* → ∞, the system follows the dynamics of a panmictic population described by equations (1 − 3). In figure 2C we used the individual-based simulations to highlight the combined effects of *ρ* and *R* on polyploid establishment.

Analysis of the system of equations (1 − 3) shows that population size is a major determinant underlying the ability of polyploids to invade a sexually reproducing diploid population. More specifically, it reduces the magnitude of meiotic errors leading to unreduced (diploid) gametes that are necessary to fully convert a population from a diploid to a tetraploid state. In space, the mating radius *R* increases drift by reducing population sizes locally, which catalyzes polyploid spread from spatial accumulation points. These observations are based on a model where sexual reproduction with allowed selfing is assumed across ploidy levels (Iterated map described by equations 1 − 3). It is however known, especially in plants, that polyploidization may lead to asexual reproduction (34, 36, 44). In this case, dispersal constraints are not expected to play a significant role in the establishment of polyploid populations since an emerging polyploid individual does not need to rely on the frequency of compatible mating partners inhabiting its surroundings, thus avoiding so-called minority cytotype exclusion. Thus, asexual reproduction has been hypothesized to facilitate polyploid establishment (34, 36). Here, we confirmed this hypothesis through spatially explicit individual-based simulations of the system defined by equations (1 − 3), assuming also strictly outcrossing and clonally reproducing polyploid individuals. For each mode of reproduction, we measured the number of generations needed for polyploids to take over the system and found that clonal reproduction allows approximately a 6-fold increase in the speed of polyploid invasion (*SI Appendix Figure S3*).

### Niche evolution and simulation of the phylogeographic history of an empirical autopolyploid complex

The deterministic and stochastic models considered thus far have demonstrated that, in the absence of fitness differences and under the assumption of sexual reproduction, polyploid establishment in space is modulated by two main ecological components: mating range (or dispersal limitation) and population density (or ecological drift). To test whether these components can explain macro ecological and evolutionary patterns in the field, we used the individual-based spatially explicit simulations with strictly outcrossing organisms which we described and analyzed earlier. However, unlike the previous models, here we also allowed the system to evolve by adding three additional components; individual fitness (independent of ploidy level), mutation rates and population growth. We started by considering a rectangular lattice of length 200 and width 100 cells. We added a linear one-dimensional environmental gradient along the length of the rectangle and started the system with an initial population size of 2000 diploid individuals adapted to the right-most corner of the landscape, but randomly distributed in the first half of the rectangle upon initialization (see *SI Appendix figure S4*). Each cell in the lattice contains a reference value (from 0 to 100), which increases linearly along the length of the rectangle from left to right, representing thus the environmental gradient, and encodes a corresponding binary sequence. The reference value denotes the percentage difference of the hamming distance between the binary sequence at the right-most corner of the lattice, i.e., start of the gradient, and the sequence at the cell where the reference value is considered. These sequences represent the optimal sub genome that leads to maximum fitness in a given location, where the fitness of an organism is a function of the smallest hamming distance found between one of its chromosomes and the encoded binary sequence. Finally, during mating, bivalent paring of chromosomes followed by a small mutation rate *μ* guarantee that offspring will inherit and evolve from parental genetical structures (see *Methods* for details).

Because selection acts across the gradient on an initially adapted diploid populations, a natural population density cline ensues. The cline thus modulates *υ*_*max*_ and, as a consequence, produces a distribution for the probability of polyploid takeover whose peak concentrates towards the edge of the ecological range occupied by diploids (figure 3A). Over the course of time, individuals start to progressively occupy the uninhabited regions of the landscape and produce ever-more adapted subpopulations. Because tetraploids emerge in those places of low-density diploid subpopulations, i.e., regions of high ecological drift, computer simulations confirmed that they are expected to emerge at the edges of niche ranges and ensure monopolization by adapting ahead of potential invading diploids (figures 3B and 3C). In this context, polyploids will monopolize ecological niches beyond the original ranges of their diploid ancestors through priority effects. In other words, accumulation of polyploidy within regions of high ecological drift – as in the edges of distribution ranges – ensure that polyploids adapt to these locations ahead of diploid ancestors and, as a consequence, are given priority for monopolization of niche expansion fronts. Niche differentiation might emerge as a consequence of a niche expansion wave that evolves over a long period of time (see *SI Appendix MovieS1* for an animated example).

**Figure 3.**
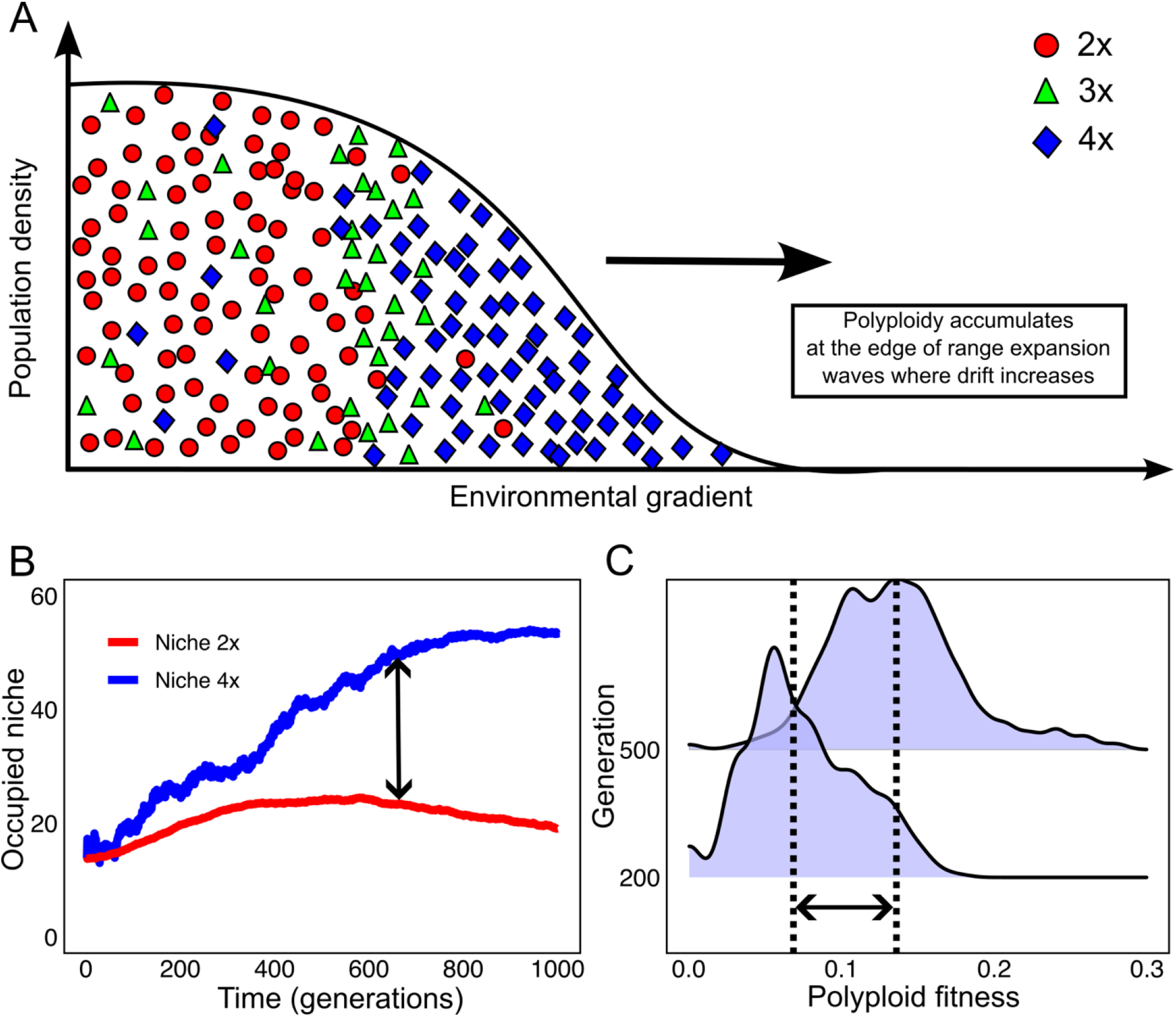
Polyploids accumulate at the edge of niche expansion waves. Panel A) is a conceptual representation of the structure of a mixed-ploidy population along a one-dimensional environmental gradient. Here, it is shown a relative higher frequency of tetraploids occurring at the edge of the natural population range. During range expansion, polyploids advance first and priority effects may ensue. In panel B) the average occupied niche (measured as the average x coordinate) factored by ploidy level is plotted as a function of time. The double arrow highlights when diploids stop advancing over the landscape which becomes increasingly monopolized by tetraploids. In C) the fitness distribution of polyploids over the second half of the landscape, i.e., for x coordinates greater than 100, is plotted for generations 200 and 500. Here, the double arrow depicts the difference in average fitness which signals adaptation. Results in B) and C) and based on individual-based and spatially explicit simulations that evolve according to the rules of inheritance and selection as described in the *Methods* section. Parameters used are *μ* = 5 × 10^−5^, *m* = 0.5, *υ* = 10^−3^, *ϕ* = 0.3 and *R* = 1.

To better illustrate the results obtained, we simulated the phylogeographic history of the *Neobatrachus* polyploid complex in Australia. The genus *Neobatrachus* is a particularly well documented polyploid complex which consists of three autotetraploid and five diploid species, whose range distributions have been shown to be well explained by average annual precipitation values across Australia (2, 4). In general, tetraploids seem to occupy the drier desertic central regions of the country, whereas diploids inhabit (mostly) the wetter regions closer to the southern coastline (2, 4). The most basal (diploid) members of the genus have an extant geographical distribution adjacent to the coastline in the southwestern region of the country (4). To simulate phylogeographic radiation within the group, we initialized simulations with a randomly distributed and well-adapted founding diploid population in the southwestern corner of simulated version of the Australian map. Instead of considering a one-dimensional linear environmental gradient on a rectangular lattice as before, we used the Australian map in conjunction with georeferenced annual precipitation (Bio12) data extracted from the *WorldClim* database (45), which was used to represent the reference values in each grid cell (see *Methods*). Thus, dispersal was constrained by real precipitation data in the model, closely approximating the modulation of range distribution in the empirical system (see *SI Appendix FigureS5* for the empirical distribution of diploids and tetraploids).

Snapshots of an instance of the simulations are shown in figure 4A (see *SI Appendix MovieS2* for an animated example). A founding diploid population progressively expands and occupies a larger portion of territory through dispersal and adaptation following a path of least evolutionary resistance. That is, dispersal and establishment are favored towards regions of similar ecological conditions. Here, polyploids will preferentially emerge at the edges of diploid populations. The same ecological components discussed previously are at play. On the one hand, reduced population densities at the leading edge of an expansion wave increase drift, which reduces *υ*_*max*_ locally and drives polyploid establishment. On the other hand, spatial clustering of tetraploid cytotypes in time ensure their persistence (figure 4B). Establishment of polyploid populations at the edges of a diploid expansion wave will block the wave itself while exposing the emerging cytotype to new ecological opportunities. The joint effects of population density and spatial dynamics lead primarily to geographical divergence which will precede any niche divergence process. Measurements of niche divergence (see *Methods*) across 1000 independent simulations demonstrated that it is a rare event with a highly positively skewed distribution. In fact, the frequency distribution of diploids and tetraploids of *Neobatrachus* species - at least along the annual precipitation gradient - is highly overlapping with shifted modes, which can be indicative of geographical clustering rather than ecological divergence (figure 4C).

**Figure 4.**
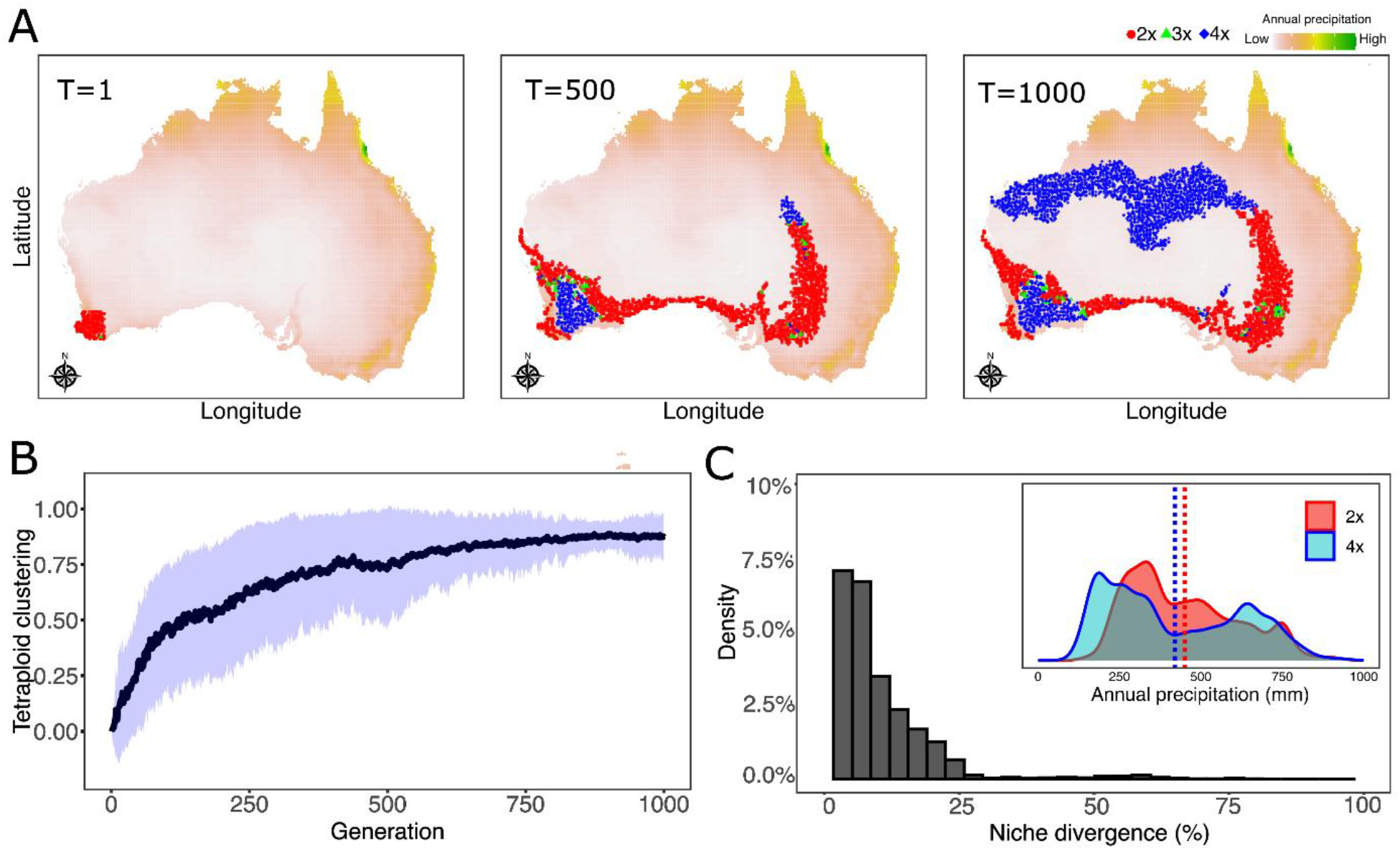
Simulated phylogeographic history of the *Neobatrachus* polyploid complex in Australia. In A) we show three snapshots of a single, independent, simulation of the phylogeographic history of *Neobatrachus* in Australia using the spatially explicit individual-based simulations described earlier. B) depicts the average tetraploid clustering (average frequency of polyploid cytotypes within the mating radius of a polyploid individual) as a function of time. In C) the frequency distribution of niche divergence (NND, see *Methods*) is shown. Inset corresponds to the empirical density distribution of *Neobatrachus* frequency – factored by ploidy level – as a function of average annual precipitation data in Australia. Vertical dotted lines correspond to average annual precipitation occupied by each ploidy level. Data from simulations are the result of 100 independent runs of the models. Average niche divergence from simulations was 8.75%. Empirical niche divergence was calculated from GBIF distribution data, https://doi.org/10.15468/dl.v6zfrw, (inset) was found to be 2.22%. Parameters used are *μ* = 5 × 10^−5^, *m* = 0.5, *υ* = 10^−3^, *ϕ* = 0.3 and *R* = 1.

## Discussion

The long-term success of related diploid and polyploid lineages depends on the magnitude of the competition caused by the overlap of their ecological niches (16). On a more elementary level, under the assumption of panmixia and only viable tetraploid and diploid cytotypes, Felber (24) and Kauai *et al*. (27) showed that the rate of the formation of unreduced (diploid) gametes, *υ*, produced by diploid cytotypes, is the only parameter controlling cytotype coexistence or polyploid invasion. In this classical model, tetraploid invasion occurs when the rate of unreduced gametes by diploid cytotypes exceeds 17.1%. Here, we found this limiting rate (*υ*_*max*_) to be 20%, by assuming that tetraploids also produce unreduced gametes at the same rate as diploids, erasing thereby a tetraploid fitness advantage. When triploids have non-zero fitness and produce haploid, diploid and triploid gametes, as observed experimentally across a wide range of taxa (27, 29), the critical *υ* was found to decay approximately exponentially as a function of triploid fitness. In other words, for each unit increase in triploid fitness, the frequency of unreduced gametes by diploid cytotypes needed for polyploids to invade the system decreases exponentially. Under small, yet non negligible, triploid fitness (*ϕ* = 0.30), *υ*_*max*_ ≈ 1.8%, which is on the same order of magnitude of frequencies found experimentally in many plant systems (30). Importantly, we showed that reduced population sizes, or increased drift, may lower *υ*_*max*_ even further. These results suggest that as triploid fitness increases across lineages, polyploid establishment and diversity must increase accordingly.

An important aspect of Felber’s model is that coexistence of cytotypes, when 0 < *υ* < *υ*_*max*_, does not represent the scenario where polyploids maintain a self-sustaining population (24). Instead, the model converges to stable cytotype frequencies that are mutually dependent. That is, if the rate of unreduced gamete formation approaches zero (*υ* → 0) at any instance of time, the existing polyploid population collapses and the system rebounds to its initial fully diploid population. Previous theoretical investigations have suggested that when mixed-ploidy populations are spatially structured, then self-sufficient polyploid populations may emerge for a more extended period of time (11, 34). Essentially, this corresponds to a scenario where the assumption of panmixia is broken. As we have shown here, polyploid spatial spread is driven by constrained mating ranges through limited dispersion. A reduced mating neighborhood directly reduces the number of available partners, which increases drift, and thus enhances the chances of polyploid takeover by locally constraining cytotype minority effects. Another aspect linked to dispersal limitation concerns the mode of polyploid invasion. Whereas in the simple case of panmictic populations polyploid takeover is abrupt past *υ*_*max*_, in space invasion proceeds iteratively by incremental coverage of the landscape. This result underlies the observation by Spoelhof *et al*. (34) that spatial structure increases the time for polyploid invasion, and is analogous to the functioning of spatial waves of advance with bistable dynamics (46).

Although in nature species coexistence is always bounded temporally (47), dispersal limitation in mixed-ploidy populations can extend the lifespan of polyploid populations. As suggested in previous work (11), constrained dispersal may increase the capacity of emerging polyploid populations to self-regulate by providing spatially concentrated subpopulations a higher frequency of compatible mating partners for a more extended period of time. While the most fundamental parameters of the mathematical model presented here, i.e., the frequency of unreduced gametes and triploid fitness, provide insights on how susceptible a sexually reproducing population is to polyploid invasion, spatial structure, which is a result of dispersal constraints, determines how stable these emerging populations become. Thus, here we identified two physiological parameters and two ecological processes that can be understood as the most elementary factors driving the establishment of polyploid populations. At a physiological level, we identify the rate of unreduced gametes and triploid fitness. Spatial sorting due to dispersal limitation and ecological drift resulting from finite population sizes underlie the dynamics of mixed-ploidy systems at an ecological level. Naturally, this result holds for polyploids that reproduce sexually with a degree of obligate outcrossing reproduction. Transition to clonal reproduction greatly facilitates polyploid invasion by overcoming the friction of constrained movement in space, as shown by our results and previous investigations in the literature (36).

Drift and spatial sorting are prominent at the margins of expanding ranges and/or after recolonization of disturbed sites (48, 49), which is in general agreement with our results. Moreover, as we have shown through simulations, a third eco-evolutionary process relevant to interploidy niche segregation is likely to arise due to monopolization effects: a species that monopolizes a novel environment first may block later arriving species by both numerical and fitness advantages (50). Such advantages are typically assigned to the ecological and/or evolutionary mediated monopolization of biotic resources. Since fitness disadvantages of cytotype hybridization decrease in a founding population of polyploids, further colonization, even by in principle fitter (diploid) cytotypes, will be blocked by the very principle that constrains polyploid establishment, i.e., minority cytotype exclusion (9). Early polyploid colonizers may evolve and adapt from mutation-generated genetic variation. Under such evolutionary mediated monopolization, adaptation augments transient numerical priority effects by accelerating early polyploid population growth, thereby promoting its long-term dominance by equalizing or elevating its fitness relative to later arriving (polyploid and non-polyploid) lineages.

To illustrate the extent to which these ecological processes can explain observed patterns empirically, we used our model to simulate the phylogeographic history of the well-documented *Neobatrachus* polyploid complex in Australia. Previous work has shown that the three autopolyploid species of the genus occupy, on average, the drier regions of the country as compared to their five diploid ancestors (2). These observations, along with other specific empirical systems, have been used to draw a correlation between polyploid niche divergence and stress (8, 19, 51, 52). Here, we have demonstrated that the frequency distribution of these populations of burrowing frogs, factored by ploidy level across the gradient of annual precipitation, is in fact highly overlapping. Polyploid frequency distributions appear bimodal with each mode displaced to one of the extremes of the distribution, away from the high concentration of diploids at the center of the gradient. Since all polyploids here are autotetraploids, this observation can be indicative of geographical displacement rather than niche differentiation. That is, emergence of polyploid populations at the edges of diploid populations might have been selected for during the establishment of the complex. The results of our simulations broadly confirm this interpretation.

The leading edge of a diploid range expansion wave contains, by definition, less dense subpopulations, which increases ecological drift and therefore reduces the limiting rate of unreduced gametes (*υ*_*max*_) locally. The spatial aggregation of polyploids in these subpopulations, through dispersal limitation, then blocks the advance of diploid cytotypes. As a result, highly concentrated populations of disparate ploidy levels occupy segregated spatial domains, which will subject these same populations to different environmental conditions and possibly very different evolutionary trajectories. Thus, our theoretical and numerical results provide evidence for the argument that the onset of polyploid niche expansion waves, or alternatively, the onset of niche divergence between ploidy levels, results from more primitive ecological processes rather than differences related to ploidy-specific genetic expression. In fact, simple models of population genetics had already suggested that there is no reason to expect that an increase in ploidy level has any bearing on adaptive potential *a priori* (27, 53). Moreover, recent empirical work shows that polyploidization leads to changes in climatic niche in an unpredictive manner (17). In other words, the magnitude and direction of niche divergence do not allow to disentangle specific ploidy-specific responses that lead to polyploid establishment. Additionally, according to our results, regardless of the ploidy level, species that occupy different spatial domains will most likely experience different selective pressures due to disparate ecological opportunities throughout their evolutionary history, which will invariably result in a myriad of niche evolution patterns.

There is accumulating evidence that polyploids emerge across the tree of life during periods of global environmental change (8, 51, 52, 54, 55). Nonetheless, the conjecture that polyploid organisms thrive in disturbed environments or have an adaptive potential, relative to their diploid ancestors, greater than expected by chance, represents a paradox. On the one hand, data show that polyploids undergo extinction rates superior to those found in diploid lineages (10, 11, 56). On the other hand, empirical work does not unequivocally support a causal correlation between adaptive responses and polyploidization in extant populations (57–59). As with polyploid niche evolution, the clustering of polyploidization events across the tree of life can be explained by simpler, more primitive, parameters and processes. For instance, at a physiological level, the rate of unreduced gametes is a fundamental driver of polyploid establishment, which has been repeatedly shown to vary substantially across taxa and as a function of environmental conditions (29, 60–62). In this context, environmental turmoil can be expected to increase production rates of unreduced gametes past *υ*_*max*_ and catalyze sustained spread of polyploids. Alternatively, under the assumption that unreduced gamete rates are unchanged with respect to environmental conditions, periods of rapid environmental change can also dramatically shrink diploid populations. Reduced population sizes increase ecological drift which constrains *υ*_*max*_ and, thus, facilitates polyploid invasion. The susceptibility of a diploid population to polyploid invasion is thus increased not by environmental boosting of the production of unreduced gametes, but rather through direct regulation of the limiting rate of unreduced gametes *υ*_*max*_. In reality, we expect that both elementary processes operate together. As a consequence, these observations offer a conservative, non-adaptationist, explanation for the clustering of polyploids during periods of environmental upheaval in the history of life.

## Methods

### General description of the modelling framework

Here, we consider an alternative description of the classical model of (auto)polyploid establishment first introduced by Felber (24). We follow previous work by us (27), where instead of tracking the time-evolution of cytotype frequencies, we model the dynamics of gamete types in a population. This alternative mathematical formulation allows one to more easily study Felber’s model analytically. An important distinction is however introduced. The model presented here also contemplates the dynamics of triploid gametes that are produced by non-segregation of chromosomes during meiosis of triploid cytotypes. This is a fundamental new property that more closely approximates empirical evidence (29). Thus, this formulation models an infinite panmictic population where diploids produce unreduced (diploid) gametes at a rate *υ* and haploid gametes at a rate (1 − *υ*). Triploid cytotypes produce haploid, diploid and triploid gametes with rates *λ*_1_, *λ*_2_ and *λ*_3_, respectively, and have a relative contribution (fitness) to the gamete pool of *ϕ* ≪ 1. As with diploids, tetraploids also produce unreduced (tetraploid) gametes with rate *υ*. We neglect tetraploid gametes since syngamy involving these gametes results in individuals with ploidy levels above five – generally found to have low fitness empirically. Individual-based simulations are used for *in-silico* experiments and to study the dynamics of Felber’s model in spatially structured populations.

### Empirically motivated eco-evolutionary simulation of mixed-ploidy populations

To simulate the dynamics of mixed-ploidy populations found empirically, we study Felber’s model on finite spatially concentrated populations. Each individual thus inherits an ordered pair of spatial coordinates (*x, y*) and possesses a set Λ_*p*_ of chromosomes with *p* elements (ploidy level). A chromosome *B*_*i*_, where *i* ∈ {1, …, *p*}, is a string of length |*B*_*i*_| = 100 with binary entries. At each generation all individuals can mate with equal probability. Mating occurs with neighbors that are at an extended Moore’s distance R from a randomly chosen seeker. A mating event involves selecting a subset of chromosomes from Λ_*p*_ (gametes) according to the rates of gamete types produced in Felber’s model. The result of the union of these subsets will make up the genome of the offspring, whose location is randomly chosen from one of the (2R + 1)^2^ cells around – and including that of – the seeker. Additionally, population size is controlled by constraining density. The number of offspring generated by each mating event is taken from a *Poisson* distribution with mean 5. Offspring dispersal is successful when the population density inside the seeker’s neighborhood is at most 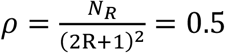, where *N*_R_ is the current number of individuals in the neighborhood of the seeker.

Movement is constrained by environmental selection. Each spatial coordinate in the landscape has an environmental value which is encoded in a string *E*_(*x,y*)_ with 100 binary entries. The encoding sequences *E*_(*x,y*)_ are built by considering an environmental gradient extracted from *WorldClim* data. For example, in this study we considered the overlap of the Australian map with *Bio12* data of 10 minutes resolution. Precipitation data rounded to one digit were then rescaled to the closed interval [1, 100] and the environmental sequences were built by randomly creating a sequence *E*^1^ for the precipitation value 100. For each unit decrease in precipitation values, then one entry *E*_(*x,y*)_ was changed keeping track of the previous changes. Thus, *E* = ⋃ *E*_(*x,y*)_, ∀ (*x, y*) is a set of sequences which can be ordered with respect to the number of changes relative to *E*^1^. Encoded sequences diverge as a function of average annual precipitation only (an in-depth explanation for sequence and map generation is given in the publicly available Java code in the GitHub repository). The probability that a seeker successfully reproduces is then 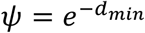, where *d*_*min*_ is the smallest Hamming distance found between one of the elements in the seeker’s genome Λ_*p*_ and *E*_(*x,y*)_ at the location of the seeker. That is, we implement chromosome-level dominance for simplicity (see results for an account of additive effects involving dominance at the allele level). Additionally, we introduce a mutation rate per nucleotide *μ* which, together with a recombination rate per nucleotide *m* during gamete formation, produces adaptive dynamics and allows populations to spread in space through evolution. Recombination is implemented with bivalent pairing of two randomly selected chromosomes during a mating event. Finally, a successful mating event happens when the Hamming distance between two randomly selected chromosomes is at least 95%, which drives speciation and spatial clustering of genetically similar individuals, mimicking macro eco-evolutionary processes (11).

To measure niche divergence between cytotypes in our simulations, we introduce a simple metric of normalized niche divergence (NND). Let 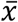 and 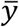 represent the average annual precipitation across tetraploid and diploid geographical locations, respectively. That is, the average environmental range occupied by each cytotype. If *e*_*min*_ and *e*_*max*_ are the minimum and maximum precipitation values found across all geographical locations, i.e., across the entire distribution range of the metapopulation, then niche divergence is calculated as:

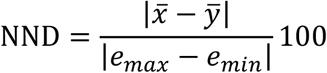

Thus, NND represents the average percentage of niche difference relative to the environmental gradient in which individuals can survive.

## Supporting information

Supplementary material - all

## Data, Materials, and Software Availability. Data, Materials, and Software Availability

Java model code and animations referred to in supplement are available at https://github.com/KauaiFe/PolyploidNiche. All additional data relevant for this study are available in *SI Appendix*.

## Acknowledgments

This work was supported by the European Research Council under the European Union’s Horizon 2020 Research and Innovation program (No. 833522) and from Ghent University (Methusalem funding, BOF.MET.2021.0005.01) (to Y.V.d.P.).

