## Supplementary material - all for "Ecological opportunity and the onset of polyploid niche expansion waves"

**This PDF file includes:**

Algorithms S1
Figures S1 to S5
Legends for movies S1 and S2

### Algorithm S1

**Algorithm S1**. Parameters: $\phi, v_{max}$

$v \leftarrow 0$ /* Initialize frequency of unreduced gametes */<br>
**while** $v < v_{max}$ **do** /* Calculate steady state frequencies up until $v_{max}$ /*<br>
    $x_t \leftarrow 1;\ y_t \leftarrow 0;\ z_t \leftarrow 0$ /* Initialize initial conditions */<br>
    converged $\leftarrow$ false /* Control whether the system has converged */<br>
    **while** converged is false **do**<br>
        /* Apply recursions from Equations 1-3 in the main manuscript */<br>
        $x_{t+1} \leftarrow Equation1(x_t, y_t, z_t)$<br>
        $y_{t+1} \leftarrow Equation2(x_t, y_t, z_t)$<br>
        $z_{t+1} \leftarrow Equation3(x_t, y_t, z_t)$<br>
    **end**<br>
    /* print cytotype frequencies for $v$ */<br>
**end**

### Supporting figures

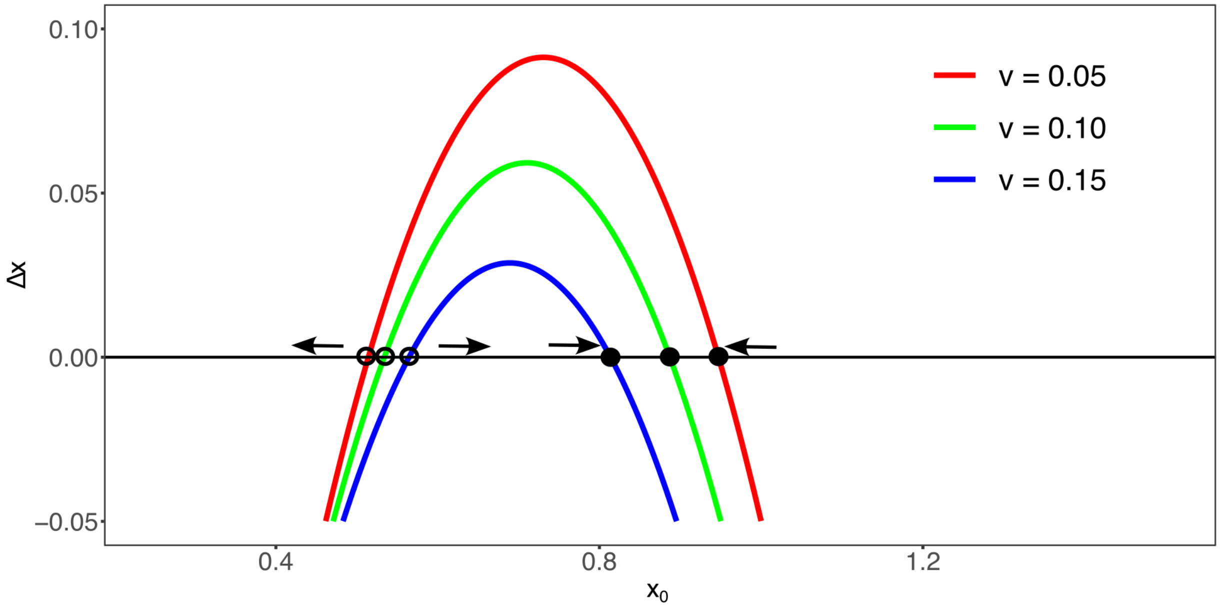


**Figure S1. Change in the frequency of haploid gametes as a function of initial conditions.** The figure depicts the change in frequency of haploid gametes (Equation 4.1 in the main manuscript) as a function of the initial frequency of haploid gametes in the system. The system is analyzed when $x_0 = 1$ because only diploids exist at time zero. Filled and unfilled circles represent stable and unstable fixed points, respectively. To illustrate stability, arrows are drawn to represent the direction of change according to the current frequency of haploid gametes. Curves are parametrized by the rate of unreduced (diploid) gametes.

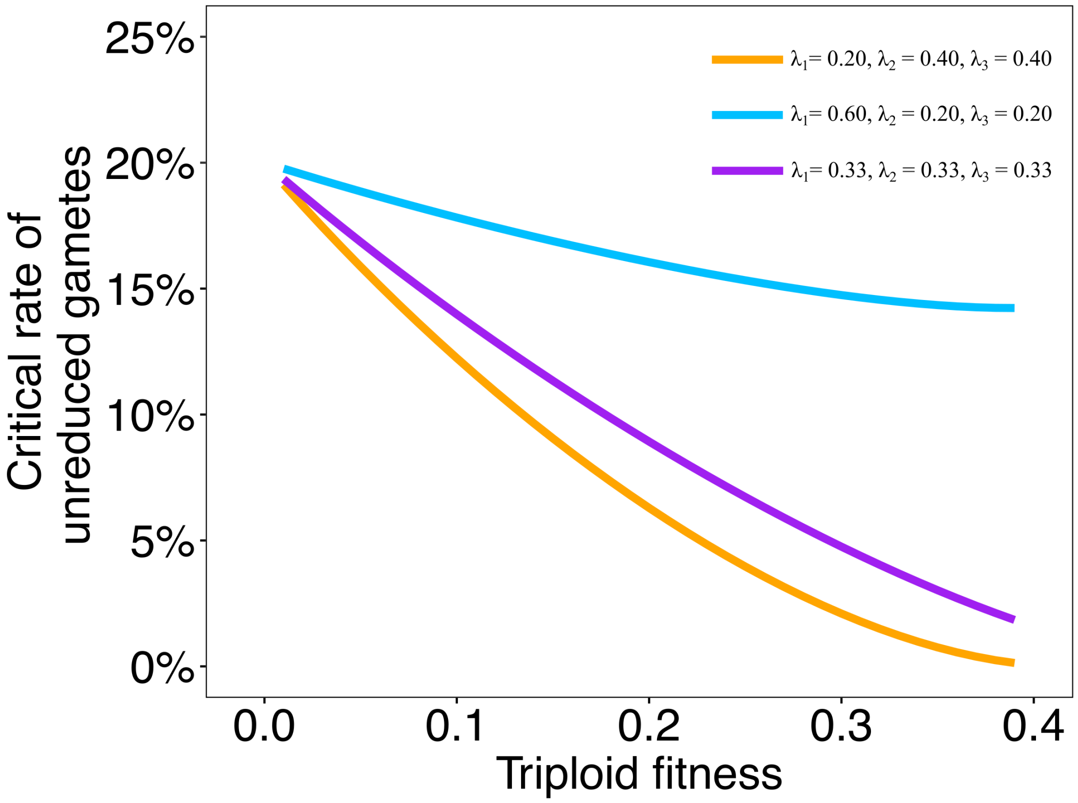


**Figure S2. Effects of the distribution of ploidy levels among gametes produced by triploid cytotypes in the susceptibility of a diploid population to polyploid invasion.** The critical rate of unreduced (diploid) gametes, $\upsilon_{max}$, is plotted as a function of triploid fitness $\phi$ (see main text). Here, $\upsilon_{max}$ was obtained heuristically for each $\phi$ in incremental steps of $0.001$. Clearly, the higher the rate of haploid gametes produced by triploid cytotypes, the greater $\upsilon_{max}$. That is, the harder it is for a diploid population to be substituted by a tetraploid one.

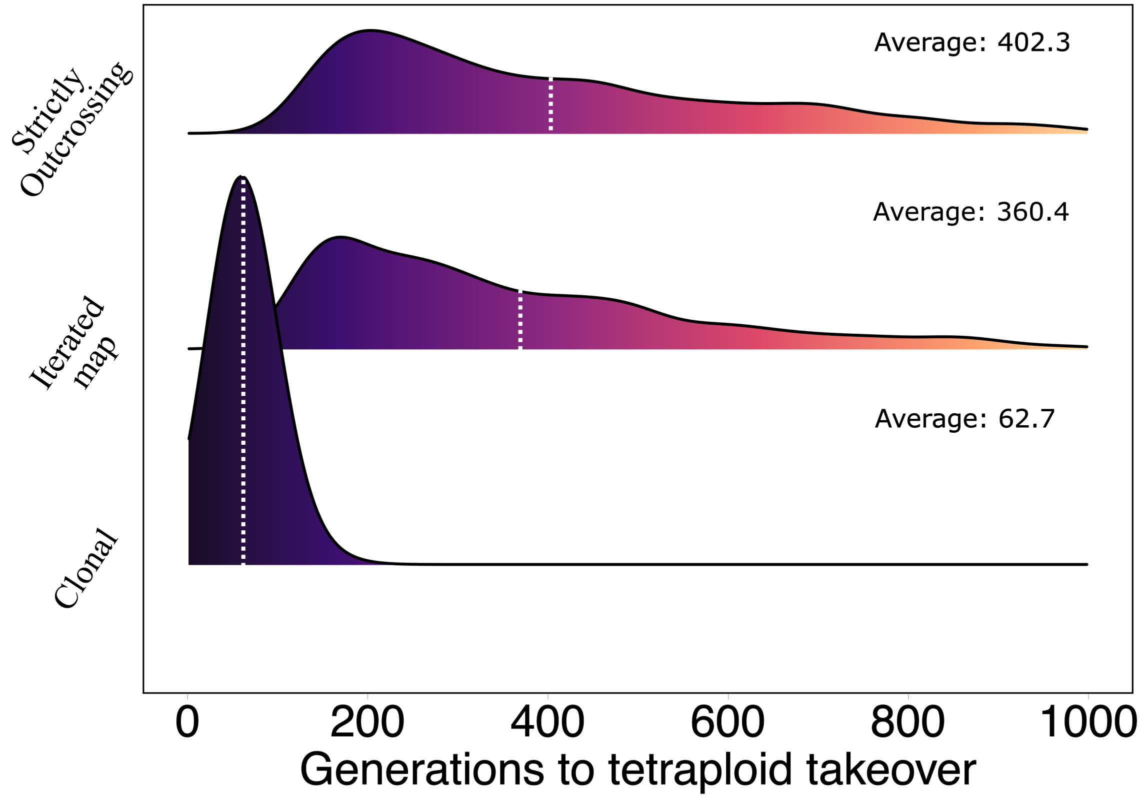


**Figure S3. Reproductive mode influences time to polyploid invasion.** The figure depicts the frequency distribution of the time required for polyploids to substitute the original diploid population in three different scenarios. These scenarios implement different modes of reproduction for tetraploids; clonal, outcrossing with selfing allowed and strictly outcrossing. Results are based on a 100 independent runs of simulations for each reproductive mode.

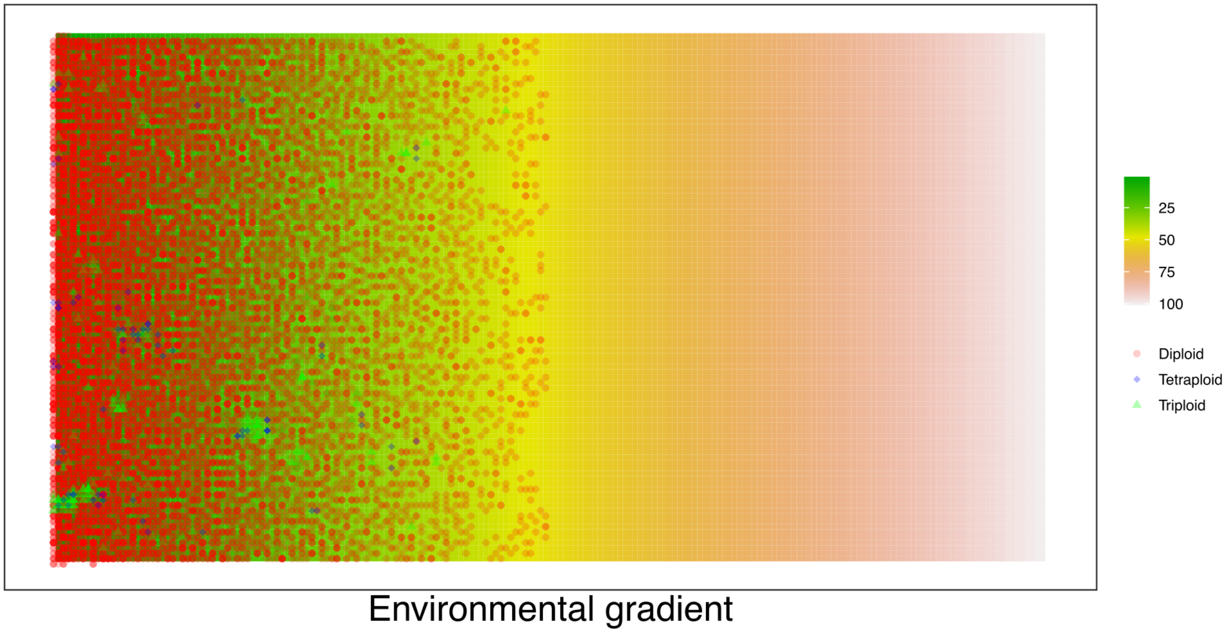


**Figure S4. Initial conditions of the individual-based spatially explicit model of polyploid establishment under selective pressure.** The figure depicts the state of an iteration of the spatially explicit model of polyploid dynamics, described within the main manuscript, at generation 5. Individuals have a genome (linear binary sequence) which is equal to the encrypted sequence at the right-most corner of the landscape (green color in the legend gradient). Due to fitness constraints for movement across the gradient, random initialization of a population guarantees a higher concentration of (fitter) individuals in the right corner of the landscape. As time unfolds, adaptive evolution guarantees progression towards the left corner.

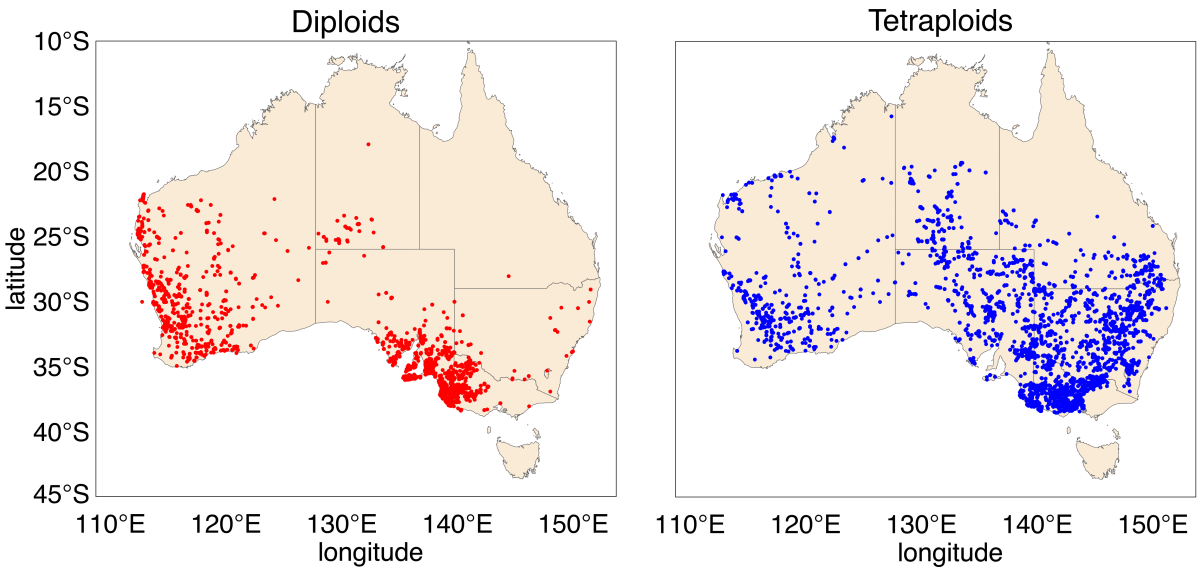


**Figure S5. Geographical distribution of diploid and tetraploid cytotypes from *Neobatrachus* in Australia.** Empirical observations of geographical distribution of *Neobatrachus* species are based on presence data from the gbif dataset: https://doi.org/10.15468/dl.v6zfrw.

#### Movies' legends

**Movie S1**. The animation depicts one instance of the individual-based spatially explicit simulation of a mixed-ploidy population evolving over a linear one-dimensional environmental gradient. Upon initialization the system is comprised of a well-adapted diploid population (red filled circles) at the left-most corner of the landscape. Dispersal, evolution and polyploidization leads to progressive range expansion towards the right where polyploids (green triploids and blue tetraploids) eventually monopolize.

**Movie S2**. The animation depicts one instance of the individual-based spatially explicit simulation of a mixed-ploidy population evolving over the Australian map. Upon initialization the system is comprised of a well-adapted diploid population at the southwestern corn of the country. Dispersal, evolution and polyploidization leads to progressive range expansion through the path of least evolutionary resistance. The accumulation of polyploidy over niche expansion fronts eventually leads to spatial segregation between cytotypes followed by adaptive responses, i.e., niche divergence.
